# Redirecting RiPP biosynthetic enzymes to proteins and backbone-modified substrates

**DOI:** 10.1101/2022.01.05.475141

**Authors:** Joshua A. Walker, Noah Hamlish, Avery Tytla, Daniel D. Brauer, Matthew B. Francis, Alanna Schepartz

## Abstract

Ribosomally synthesized and post-translationally modified peptides (RiPPs) are peptide-derived natural products that include the FDA-approved analgesic ziconotide^1,2^ as well as compounds with potent antibiotic, antiviral, and anticancer properties.^3^ RiPP enzymes known as cyclodehydratases and dehydrogenases represent an exceptionally well-studied enzyme class.^3^ These enzymes work together to catalyze intramolecular, interresidue condensation^3,4^ and aromatization reactions that install oxazoline/oxazole and thiazoline/thiazole heterocycles within ribosomally produced polypeptide chains. Here we show that the previously reported enzymes MicD-F and ArtGox accept backbone-modified monomers, including aramids and beta-amino acids, within leader-free polypeptides, even at positions immediately preceding or following the site of cyclization/dehydrogenation. The products are sequence-defined chemical polymers with multiple, diverse, non-alpha-amino acid subunits. We show further that MicD-F and ArtGox can install heterocyclic backbones within protein loops and linkers without disrupting the native tertiary fold. Calculations reveal the extent to which these heterocycles restrict conformational space; they also eliminate a peptide bond. Both features could improve the stability or add function to linker sequences now commonplace in emerging biotherapeutics. Moreover, as thiazoles and thiazoline heterocycles are replete in natural products,^5–7^ small molecule drugs,^8,9^ and peptide-mimetic therapeutics,^10^ their installation in protein-based biotherapeutics could improve or augment performance, activity, stability, and/or selectivity. This work represents a general strategy to expand the chemical diversity of the proteome beyond and in synergy with what can now be accomplished by expanding the genetic code.

## Introduction

Ribosomally synthesized and post-translationally modified peptides (RiPPs) are peptide-derived natural products that include the FDA-approved analgesic ziconotide^1,2^ as well as compounds with potent antibiotic, antiviral, and anticancer properties.^3^ RIPP biosynthesis begins with a ribosomally synthesized polypeptide whose N-terminal leader sequence (∼20 - 110 aa) recruits one or more endogenous enzymes capable of diverse post-translational modification (PTM) of an adjacent C-terminal substrate sequence.^3,11^ Researchers have leveraged this leader-dependent mechanism to direct RiPP PTM enzymes to C-terminal substrate sequences containing diverse non-canonical α-amino acids (nc-α-AAs).^12,13^

Cyclodehydratases and dehydrogenases represent an exceptionally well-studied class of RiPP enzymes.^3^ These enzymes work together to catalyze intramolecular cyclization^3,4^ and subsequent aromatization reactions that install oxazoline/oxazole and thiazoline/thiazole heterocycles within polypeptide chains (**Figure 1A, B**). Previous work has shown that the cyclodehydratases PatD^14^ and TruD^15^ support leader sequence-dependent oxazoline/thiazoline formation within substrates containing nc-α-AAs adjacent to^16^ or at the cyclization site itself.^17–19^ In related work, it was shown that a chimeric leader peptide could direct the cyclodehydratase LynD^15^ and the dehydrogenase TbtE^20^ to install thiazol(in)es within substrates containing nc-α-AAs adjacent to the cyclization site.^21^ Finally, reconstituted lactazole biosynthesis,^22^ including the cyclodehydratase-dehydrogenase pair LazDE/LazF, was found to install oxazoles and thiazoles within polypeptide substrates containing α-hydroxy, *N*-methyl, cyclic α-, and β^3^-amino acids^23^ at sites distal from the site of heterocyclization (> 4 residues away).

**Figure 1.**
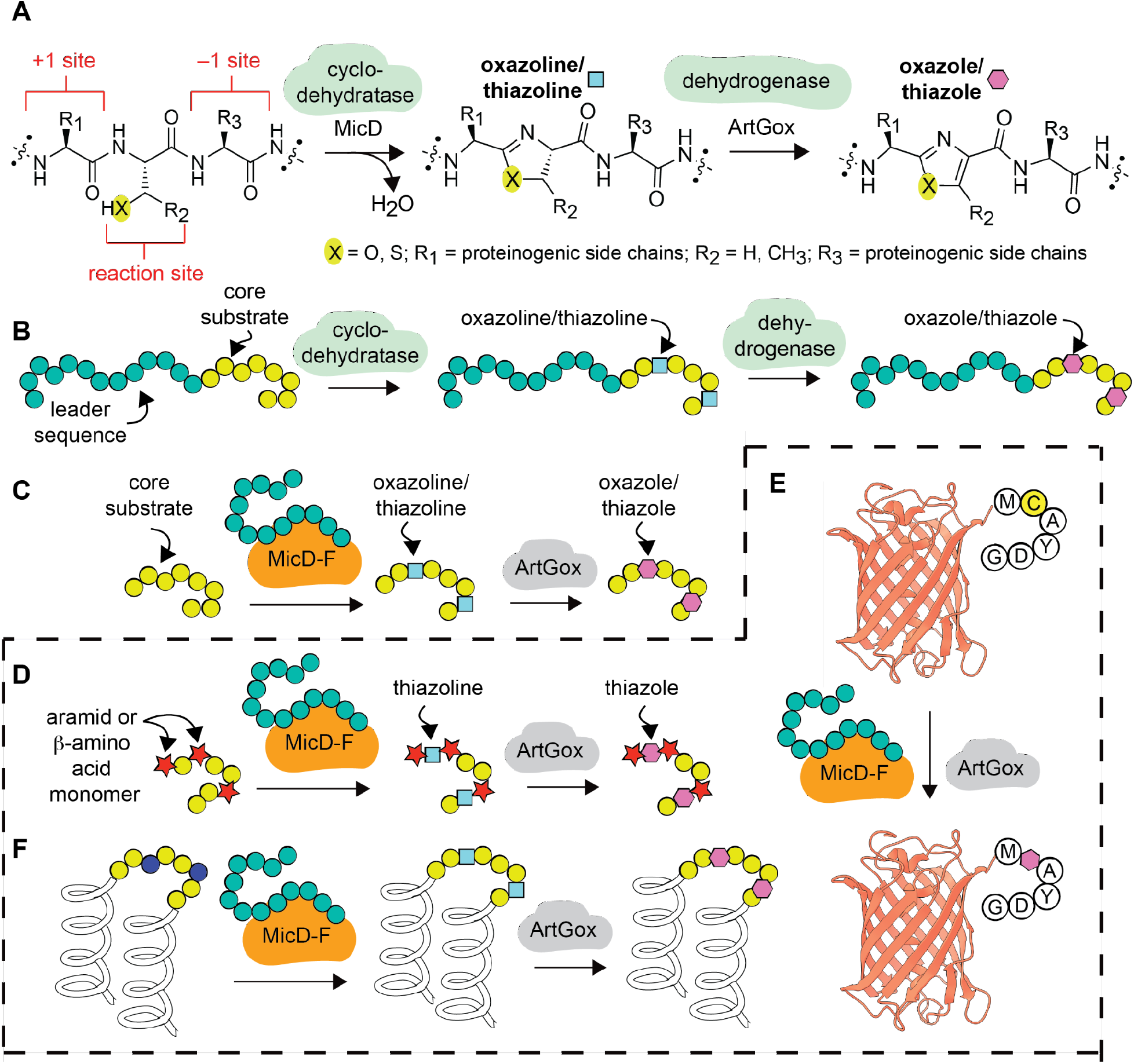
Overview of cyclodehydratase/dehydrogenase chemistry. **(A)** Scheme illustrating the natural conversion of a serine, threonine, or cysteine-containing polypeptide into a oxazoline or thiazoline-containing product through the action of MicD^26^ and subsequent dehydrogenation into an oxazole or thiazole through the action of ArtGox.^29^ (B) Natural substrates for MicD^37^ and ArtGox^38^ consist of a core sequence that includes the reaction site and an extended upstream leader sequence. (C) Fusion of the leader sequence to the N-terminus of MicD generates a constitutively activated enzyme MicD-F that processes leaderless substrates.^26^ (D) **This work**: MicD-F and ArtGox accept polypeptide substrates containing diverse non-α-amino acid monomers, including aramids, at the +1 and -1 site. MicD-F and ArtGox can also install thiazoline and thiazole linkages within globular proteins such as (E) mCherry and (F) Rop.

Previous work has also shown that certain cyclodehydratase enzymes retain activity for leader sequence-free substrates when the leader peptide is provided in *trans* (**Figure 1C**).^24,25^ Building on this observation, Naismith and coworkers engineered a family of cyclodehydratases in which the leader peptide is fused to the N-terminus of the cyclodehydratase catalyst as opposed to the N-terminus of the substrate polypeptide. These constitutively activated enzymes, notably LynD Fusion (LynD-F)^25^ and MicD Fusion (MicD-F)^26^ act in a leader peptide-independent manner to promote the cyclodehydration of polypeptides containing a C-terminal Ala-Tyr-Asp (AYD) recognition sequence.^4,25–28^ In complementary work, Schmidt and coworkers demonstrated that two dehydrogenases, ArtGox and ThcOx, also accept leaderless peptide substrates.^29^ Taken together, these enzymes represent a fully leader-free route towards polypeptides (and proteins, *vide infra*) containing mRNA-programmed thiazole and oxazole linkages. Indeed, some tolerance for non-canonical α-amino acid residues has been reported: LynD-F was shown to install a thiazoline in a peptide substrate containing 3-azido-L-alanine positioned 4 residues away from the site of cyclization,^28^ and the combination of LynD-F and ArtGox installed a thiazole in an AYD-containing peptide with a polyethylene glycol spacer 2 residues from the site of cyclization.^27^

Here we report that MicD-F^26,29^ and ArtGox^26,29^ act together to process polypeptide substrates containing diverse translation-compatible^30–35^ aramid and β-amino acid monomers, even at sites directly flanking the reaction site (**Figure 1D)**. We show further that MicD-F^26,29^ and ArtGox^26,29^ process substrates even when the CAYD sequence is positioned at the C-terminus of mCherry, a large β-barrel protein, or embedded within the loop of the dimeric α-helical bundle protein Rop; the products are folded, globular proteins containing a conformationally restricted, fully unnatural, heterocyclic backbone. To the best of our knowledge, these studies represent the first example of leader-free azol(in)e biosynthesis within polypeptides containing diverse non-α-amino acid monomers flanking the site of cyclization, and the first report of a cooperatively folded protein containing a post-translationally installed heterocyclic ring.^36^ The effects of the embedded heterocycle on local conformational flexibility are examined computationally, providing important insight into the backbone restrictions that could be leveraged to improve the physio-chemical properties of therapeutic proteins. This work represents a general strategy to expand the chemical diversity of the proteome beyond and in synergy with what can now be accomplished by expanding the genetic code.

## Results

### MicD-F and ArtGox accept substrates with diverse structures at the +1 site

We began by exploring the tolerance of MicD-F for sequences containing non-α-amino acid monomers at the +1 site (**Figure 1A** and **Figure 2A-B**). A series of nine potential substrates were prepared in which a non-α-amino acid preceded the reaction site (substrates 1(a-i)). Monomers evaluated included arenes, aramids, fluorophores, and linear and cyclic β-amino acids. All substrates contained a C-terminal AYD recognition sequence^39^ and were incubated with MicD-F (**Supplementary Figure 1A-B**) (5 mol%) under mild conditions (pH 8.0, 37 °C, 4 h) that resulted in complete conversion of ICAYDG, a substrate with the natural α-amino acid Ile at the +1 site (**Supplementary Figure 2**). Cyclization was analyzed initially via liquid chromatography-mass spectrometry (LC-MS) and the extent of product formation estimated by integrating the extracted ion chromatogram (**Figure 2B-C**). Virtually every substrate examined underwent MicD-F-catalyzed cyclization to the corresponding thiazoline under these conditions. Substrates containing electron-withdrawing or donating aromatic rings, bulky multi-ring systems, and linear and cyclic β-amino acids were all cyclized efficiently by MicD-F, with yields between 91 and 99% (products 2(a-i)). UHPLC analysis of each reaction mixture confirmed that thiazolines 2a-i were the sole reaction products under these conditions (**Supplementary Figure 3**). It is notable that monomers with highly divergent structures are accepted almost equally by MicD-F, suggesting that the +1 residue interacts minimally if at all with the enzyme active site.

**Figure 2.**
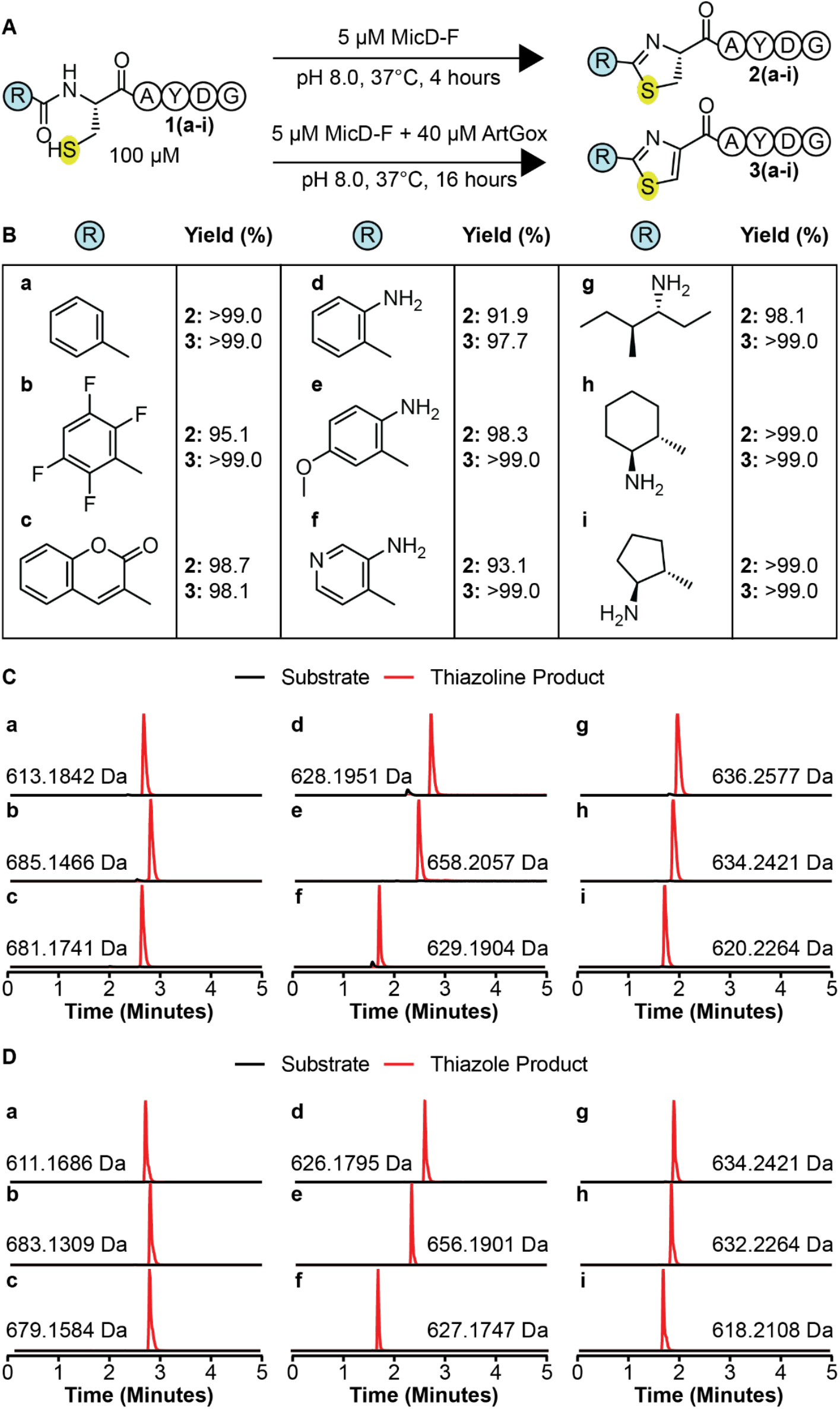
MicD-F and ArtGox tolerate diverse non- proteinogenic, non-α- amino acid monomers at the +1 site. (A) Scheme illustrating the conditions used for the reaction of ArtGox and/or MicD-F with substrates containing non-α-amino acid monomers N-terminal to the reaction site (+1 site). (B) Yields of thiazoline 2(a-i) and thiazole (3(a-i)) products obtained for substrates containing non-α-amino acid monomers at +1 site. Extracted ion chromatograms illustrating the products of (C) MicD-F and (D) MicD-F + ArtGox-catalyzed reactions.

Next, we explored whether MicD-F and ArtGox could act in synergy to convert peptides containing non-α-amino acids at the +1 site directly into the corresponding thiazoles 3(a-i) (**Figure 2A**). Substrates 1(a - i) were incubated with MicD-F (5 mol%) and ArtGox (**Supplementary Figure 1A and C)** (40 mol%) under conditions (pH 8.0, 37 °C, 16 h) that resulted in complete two-step conversion of ICAYDG into the corresponding thiazole product (**Supplementary Figure 4**). ArtGox efficiently oxidized each thiazoline to the corresponding thiazole in yields that exceeded 97% over the two steps for every example (products 3(a-i)) (**Figure 2B and D**). UHPLC analysis of each reaction mixture confirmed that thiazoles 3a-e were the sole reaction product. Coelution with excess flavin mononucleotide precluded UHPLC analysis of thiazoles 3f-i (**Supplementary Figure 5**). These results indicate that MicD-F and ArtGox tolerate diverse non-proteinogenic, non-α-amino acid monomers at the +1 site. Many of these non-α-amino acid monomers have been installed at the N-termini of ribosomally translated peptides *in vitro*,^33,34,40,41^ suggesting a path towards proteins and polypeptides with highly unique N-terminal appendages.

### MicD-F and ArtGox accept substrates with diverse structures at the -1 site

Next we explored whether MicD-F and ArtGox would accept leader-free polypeptide substrates containing non-α-amino acid monomers at the -1 site (**Figure 3**). We explored a diverse array of monomers – β^3^-amino acids, β^2^-amino acids, cyclic β^2^,β^3^-amino acids, as well as substituted and unsubstituted aramids. Notably, it was found that inserting an α-amino acid (Ile) residue between the site of cyclization and the C-terminal AYD motif necessitated higher concentrations (50 mol%) of MicD-F and up to 24 h reaction time to complete the cyclodehydration reaction (**Supplementary Figure 6**).

**Figure 3.**
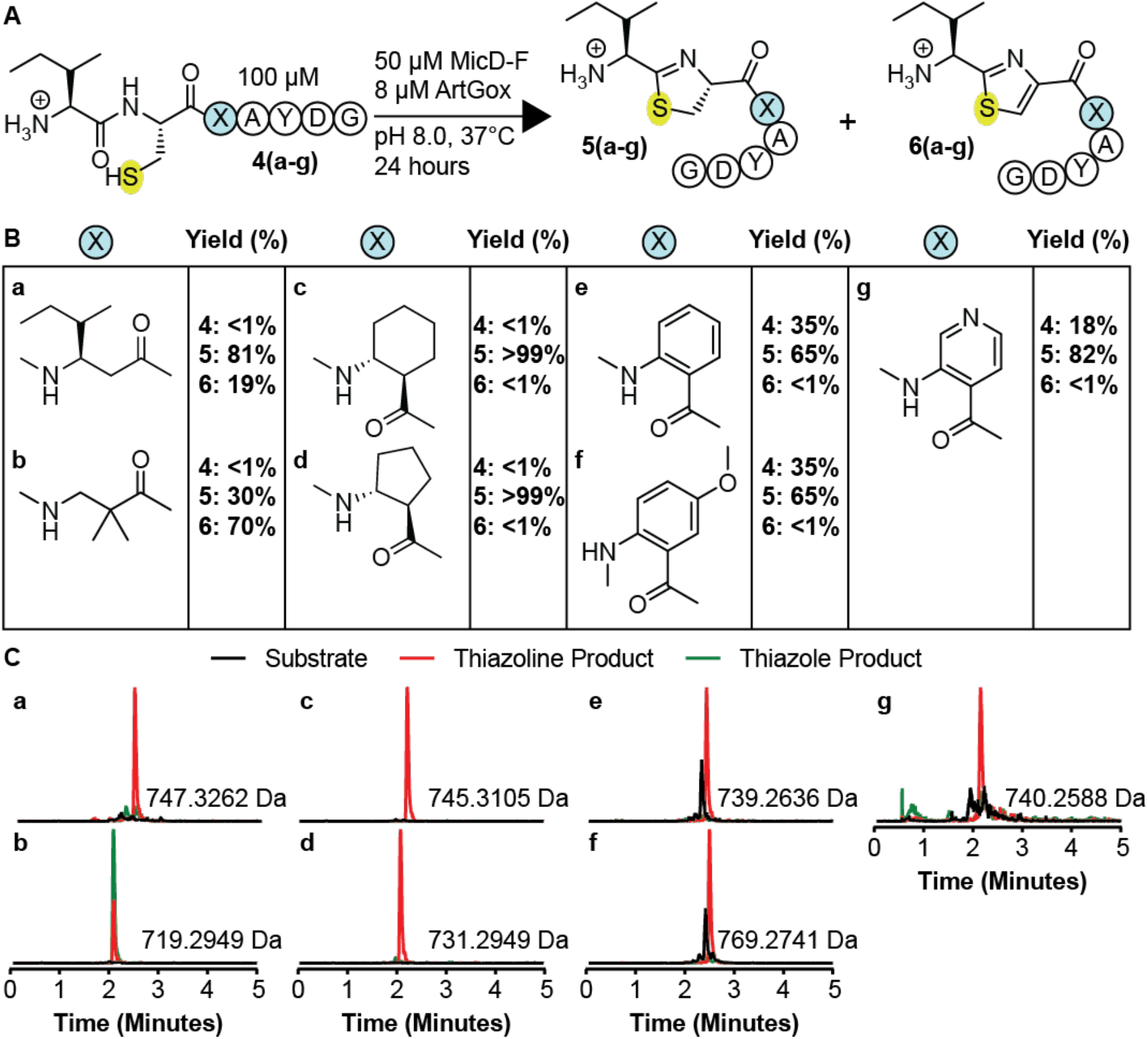
MicD-F and ArtGox tolerate diverse non-proteinogenic, non-α-amino acid monomers at the -1 site. (A) MicD-F and ArtGox reactions of substrates with non-α-amino acid monomers immediately C-terminal to the site of cyclization. (B) Yields of thiazoline 5(a-i) and thiazole (6(a-i)) products obtained for polypeptide substrates with non-α-amino acid monomers at position -1, immediately C-terminal to the reaction site. Extracted ion chromatograms illustrating the products of (C) MicD-F and (D) ArtGox-catalyzed reactions.

All -1 site substrates (substrates 4(a-g), **Figure 3A-B**) contained a C-terminal AYD recognition sequence^39^ and were incubated with MicD-F (50 mol%) and ArtGox (8 mol%) under conditions (pH 8.0, 37 °C, 24 h) that resulted in complete conversion of a substrate with a natural α-amino acid at the -1 site to the corresponding thiazole (**Supplementary Figure 7**). Reactions were analyzed as described above. Under these conditions, every peptide evaluated was a substrate for MicD-F and a few were substrates for both MicD-F and ArtGox (**Figure 3B-C**). Substrates containing β^3^-amino acid-, β^2^-amino acid-, or cyclic β^2^,β^3^-amino acids at the -1 site (substrates 4(a-d)), were fully consumed under these conditions (<1% unmodified peptide). Those with β^3^-alkyl substituents (4a, c, and d) were converted cleanly into the corresponding thiazolines 5a, c, and d, with little (4a) or no (4c,d) thiazole formation. In contrast, substrate 4b, with geminal β^2^-methyl substituents, was converted into a 30/70 mixture of thiazoline 5b and thiazole 6b. Substrates 4e-g containing aramid monomers at the -1 position reacted more slowly under these conditions, producing the analogous thiazoline products in 65-85% yield after 24 h reaction at pH 8 (**Figure 3B-C**). Surprisingly, while all substrates containing +1 site modifications were efficiently oxidized to the corresponding thiazole (**Figure 2B and D**), only the substrate containing a β^2^-amino acid modification at the -1 site was efficiently oxidized by ArtGox (70%) (**Figure 3B-C**). With the exception of substrate 4c, increasing the pH to 9.0 promoted formation of the desired thiazole product (**Supplementary Figure 8B-C**). However, even under these conditions only substrate 4b (88%) yielded greater than 41% thiazole product (**Supplementary Figure 8B-C**). These data indicate that MicD-F and ArtGox are both less tolerant of non-α-amino acid monomers at the -1 site than at the +1 site. ArtGox appears especially intolerant of substitution or sp^2^ hybridization at the β^3^-position of substrates at the -1 site.

### MicD-F is sensitive to amino acid identity at the cyclization site

To complete the exploration of the substrate tolerance of MicD-F and ArtGox, we synthesized a set of potential substrates containing a non-α-amino acid directly at the cyclization site. Each contained a C-terminal AYD sequence preceded by either L-β^3^- or D-β^3^-threonine (**Supplementary Figure 9A**). Incubation of these substrates with MicD-F (50 mol%) under conditions (pH 8.0, 37 °C, 24 h) that resulted in substantial cyclization of a substrate containing L-α-threonine at the cyclization site led to no detectable cyclization (<1%) (**Supplementary Figure 9B**). Even at pH 9.0 no cyclization occurred (**Supplementary Figure 9C**), indicating that MicD-F is highly sensitive to amino acid identity at the site of cyclization. This result is in line with previous work that demonstrated the cyclodehydratase PatD failed to react with substrates containing D-α-threonine at the cyclization site.^18^

### Redirecting RiPP biosynthetic enzymes to intact folded proteins

Thiazolines and thiazole are replete in natural products^42–44^ and synthetic drug-like small molecules,^8,9^ and calculations confirm the expected decrease in conformational freedom that derives from aromatic and/or sp^2^ character within the peptide backbone.^45^ This finding and the leader-independent nature of MicD-F and ArtGox-mediated thiazol(in)e biosynthesis inspired us to explore substrates in which the site of cyclodehydration/dehydrogenation is embedded within a stable protein fold (**Figure 4**). We first asked whether MicD-F and ArtGox could install thiazol(in)e linkages within loops and/or at the termini of mCherry. mCherry is a prototypic fluorescent beta-barrel protein derived from DsRed, isolated originally from Discosoma sea anemones.^46^ We cloned, expressed, and purified a set of mCherry variants in which the core sequence MCAYDG was appended to the mCherry C-terminus (mCherryC+) or inserted into a loop immediately downstream of residues D137 (mCherry137+), D174 (mCherry174+), V192 (mCherry192+), or E211 (mCherry211+) (**Figure 4A, Supplementary Table 2, Supplementary Figure 10**). Although mCherry137+ and mCherry211+ were partially/completely non-fluorescent or could not be purified, mCherryC+, mCherry174+ and mCherry192+ were soluble and fluorescent. In all three of these cases, mass spectrometry of the purified proteins showed the characteristic loss of 22 Da indicating chromophore maturation (**Supplementary Figure 11**).

**Figure 4.**
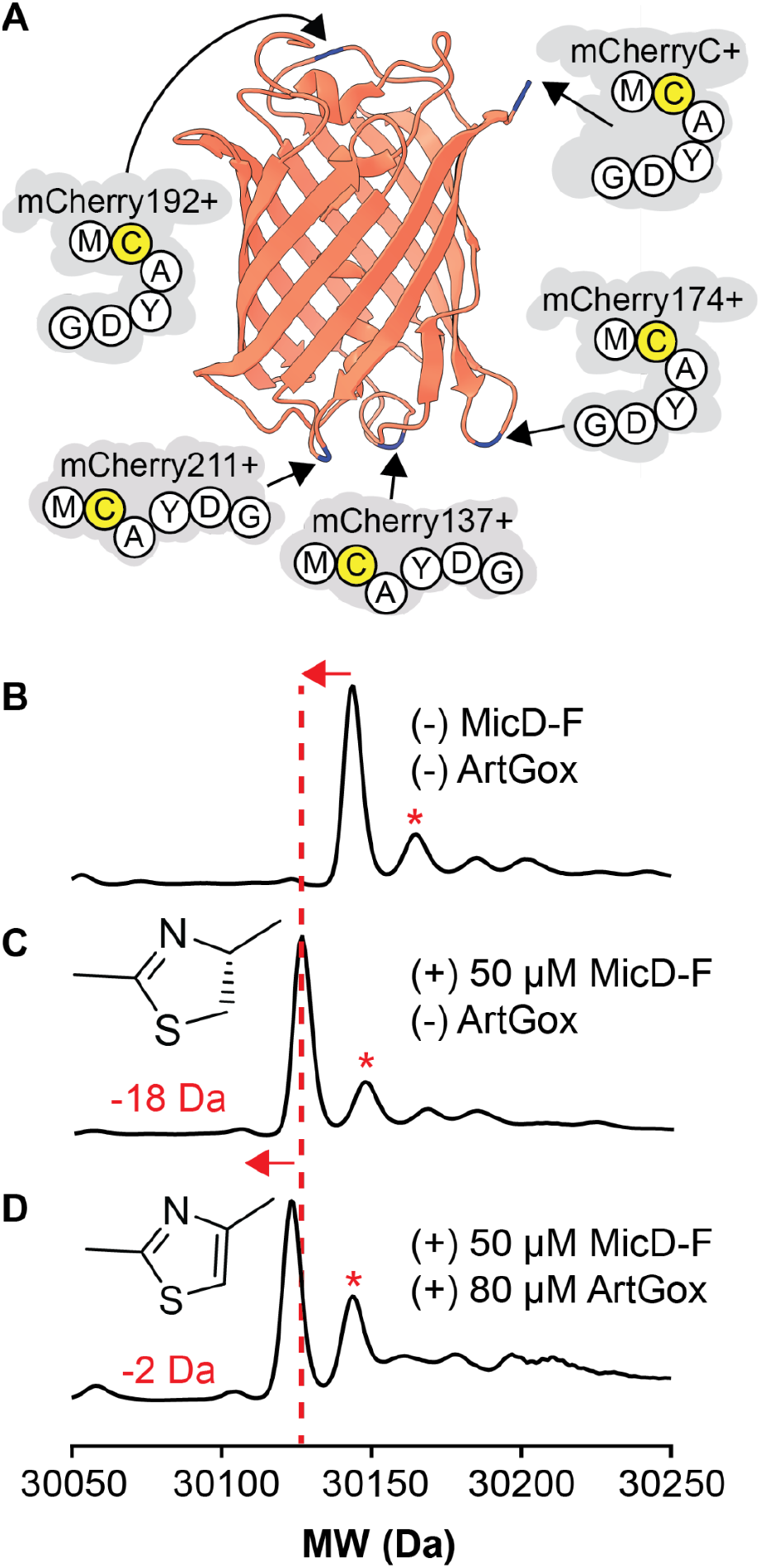
(A) mCherry variants evaluated as substrates for ArtGox and/or MicD-F.Each variant contains the sequence MCAYDG inserted following the residue shown. LC/MS analysis of mCherryC+ both (B) before and (C) after reaction with 50 mol% MicD-F or (D) 50 mol% MicD-F and 80 mol% ArtGox. Data reported are normalized counts from deconvoluted mass spectra. Asterisk indicates the molecular weight of the parent protein without a mature chromophore (+22 Da).

Treatment of mCherryC+ with 50 mol% MicD-F (pH 9.0, 24 hours, 37°C) led to virtually complete conversion to the thiazoline product as indicated by a loss of water in the deconvoluted mass spectrum (**Figure 4B** and **C**). No such mass change was observed in an analogous reaction containing mCherryC-, which carries the sequence MAAYDG in place of MCAYDG at the C-terminus, providing evidence that the observed cyclodehydration demanded a Cys residue immediately upstream of the AYD recognition sequence (**Supplementary Figure 12B-C**). Neither mCherry174+ nor mCherry192+ displayed the loss of water characteristic of successful cyclodehydration even after 24 hours at 37°C (**Supplementary Figure 12D-E**). Nevertheless, we explored the potential for MicD-F and ArtGox to act in tandem to install an aromatic thiazole backbone in mCherryC+. Simultaneous treatment of mCherryC+ for 24 hours (pH 9.0, 37°C) with MicD-F (50 mol%) and ArtGox (80 mol%) resulted in the expected -2 Da shift in the deconvoluted mass spectrum (**Figure 4D**) relative to that of mCherryC+ treated with only MicD-F (**Figure 4C**). This result indicates that the MicD-F/ArGox enzyme pair can post-translationally install an aromatic thiazole backbone within a structurally unconstrained region of a well-folded beta-barrel protein.

We hypothesized that the absence of cyclodehydration reactivity for mCherry174+ and mCherry192+ at 37°C was due to neighboring structural elements that disfavor productive interaction with MicD-F and/or enzyme-promoted thiazoline formation. Therefore, we carried out a second set of cyclodehydration reactions at 42°C, the highest temperature at which MicD-F remained stable in our hands, which should increase the conformational flexibility of loop insertions. At this elevated temperature, mCherryC+ again displayed cysteine-specific loss of water characteristic of successful cyclodehydration (**Supplementary Figure 13B-C**). However, again neither mCherry174+ or mCherry192+ displayed the loss of water characteristic of successful cyclodehydration after 24 hours at 42°C (**Supplementary Figure 13D-E**). It has been reported^47^ that the apparent melting temperature of mCherry is upwards of 90°C. Taken together with our results, this finding suggests that there is an inherent mismatch between the temperature stability of MicD-F and the thermodynamic stabilities of the mCherry loop insertions evaluated here.

To test this hypothesis, we sought a folded, globular protein with a lower melting temperature than mCherry with the expectation that it would be more amenable to insertion of an internal thiazol(in)e linkage. Rop is a homodimeric four-helix bundle protein formed by the antiparallel association of two helix-turn-helix monomers.^48^ Regan and coworkers reported many years ago that the native two-residue turn in Rop could be replaced by up to ten glycine residues without loss of the native dimer structure. The Rop variant with the longest insertion–Gly_10_–melted cooperatively at 50°C,^49^ suggesting that it might tolerate an internal, intra-loop thiazole or thiazoline (**Figure 5A**). To test this hypothesis, we expressed and purified three Rop variants containing a single CAYD sequence embedded near the N-terminus (RopN), the C-terminus (RopC), or centrally (RopM) within a ten-residue glycine-rich loop (**Figure 5A, Supplementary Table 3, Supplementary Figure 14**, and **Supplementary Figure 15**). All three Rop variants exhibited high α-helical content at 20 μM as judged by wavelength-dependent CD measurements (**Figure 5B)**. RopC and RopM migrated as discrete dimers at 50 μM as judged by size-exclusion chromatography (SEC) and melted cooperatively and reversibly with T_M_ values of 28°C and 32°C (**Supplementary Figure 16)**. RopN, by contrast, migrated as a heterogeneous mixture upon SEC and melted non-cooperatively, albeit at a slightly higher apparent T_M_ (43°C) perhaps because of disulfide formation (**Supplementary Figure 16)**.^50^

**Figure 5.**
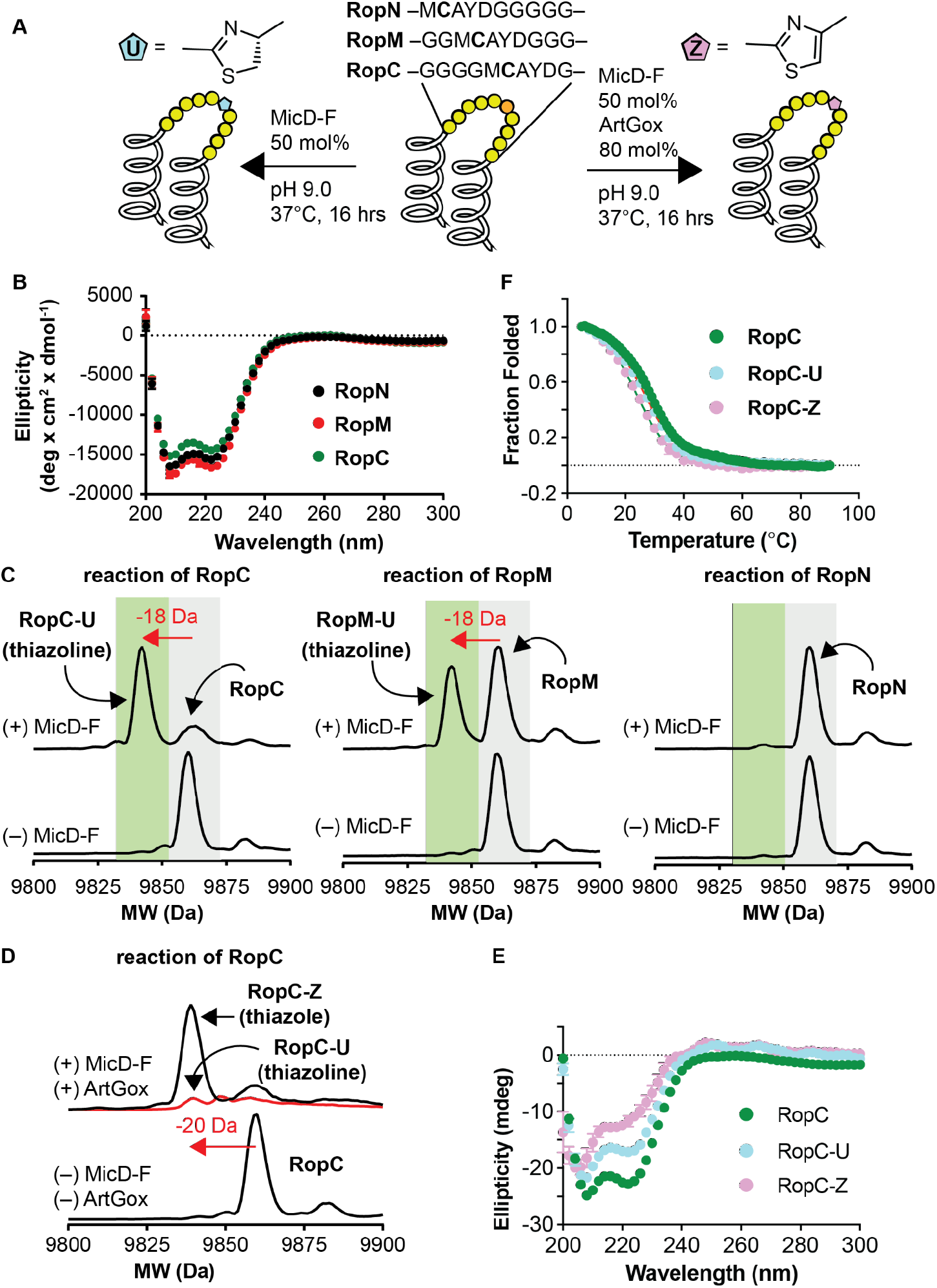
MicD-F and ArtGox act in tandem to install thiazoline and thiazole backbones within globular proteins. (A) Cartoons illustrating the sequences of RopC, RopM, and RopN and the conditions used for MicD-F-catalyzed cyclodehydration (left arrow) or tandem cyclodehydration/dehydrogenation catalyzed by MicD-F and ArtGox (right arrow). (B) Wavelength-dependent circular dichroism spectra of RopC, RopM, and RopN at [monomer] = 20 μM in 10 mM phosphate, 100 mM NaCl, 150 μM TCEP, pH 7.0 and 25°C. (C) LC/MS analysis of the reaction of RopC, RopN, and RopM with MicD-F under the conditions shown in panel (A) above. The characteristic loss of 18 mass units upon cyclodehydration is evident for both RopC and RopM; the reaction of RopM was incomplete under these conditions. (D) Treatment of RopC with MicD-F and ArtGox under the conditions shown in panel (A) above led to clean conversion into the corresponding thiazole (RopC-Z). (E) The wavelength-dependent CD spectra of RopC-U and RopC-Z compared to RopC; these data are not corrected for contributions due to the thiazoline or thiazole linkage. (F) The melting temperatures of RopC, RopC-U, and RopC-Z (after refolding) are almost identical.

Although RopC, RopN, and RopM all contained the same CAYD recognition sequence, only one–RopC–underwent clean conversion into the corresponding thiazoline upon treatment with 50 mol% MicD-F (pH 9.0, 37°C, 16h). RopM reacted partially under these conditions and RopN was unreactive (**Figure 5C**). Reaction of RopC to generate thiazoline RopC-U proceeded more slowly at 25 °C (**Supplementary Figure 17**). RopC could be converted directly into the thiazole RopC-Z upon treatment with 50 mol% MicD-F and 80 mol% ArtGox (**Figure 5D, Supplementary Figure 18**). No reaction was observed when the Cys residue within the RopC reaction site was replaced with Ala or when the C-terminal AYD sequence was replaced by GGG **(Supplementary Figure 19)**.

The products of the reaction of RopC with MicD-F (RopC-U) and with MicD-F and ArtGox (RopC-Z) were purified and analyzed by size-exclusion chromatography and wavelength- and temperature-dependent CD. Thiazoline-containing RopC-U was a homogeneous dimer as judged by SEC (**Supplementary Figure 16)** and retained a significant level of α-helical structure (**Figure 5E**). It also melted cooperatively and reversibly with a T_M_ value of 27 °C, a value almost identical to that of RopC itself (28 °C) (**Figure 5F**). Thiazole-containing RopC-Z displayed more complex behavior; it was less homogeneous as judged by SEC and melted cooperatively (T_M_ = 24 °C) but only after a refolding step (**Supplementary Figure 20**). These results indicate that the MicD-F can post-translationally install a thiazoline within a backbone of a helical bundle protein, and that ArtGox can oxidize this substrate to install a fully aromatic thiazole unit.

### Computational analysis of the effects of thiazoline/thiazole formation on local backbone flexibility

To explore the effects of the cyclization reaction on local backbone flexibility, we examined the conformational space of the tetrapeptide Ac-AACA-NH_2_. The use of this simplified substrate allowed the inherent energetics of the backbone to be evaluated in the protein context without the complications of side chain fluctuations. Molecular mechanics methods (Macromodel, OPLS4 force field, implemented in Schrödinger Maestro software) were first used to generate and minimize large populations of conformers for cysteine-, thiazoline-, and thiazole-containing analogs (**Figure 6A**). For each species, 10,000 starting structures were sampled using the Mixed Torsional/Low-Mode method. All conformers within 4 kcal/mol of each global minimum were then subjected to geometry optimization using DFT (Jaguar: B3LYP-D3/6-31G**). An SM8 method was used to determine the relative energies in aqueous media.^51^ All non-redundant conformers were then ranked based on these energies and compared.

**Figure 6.**
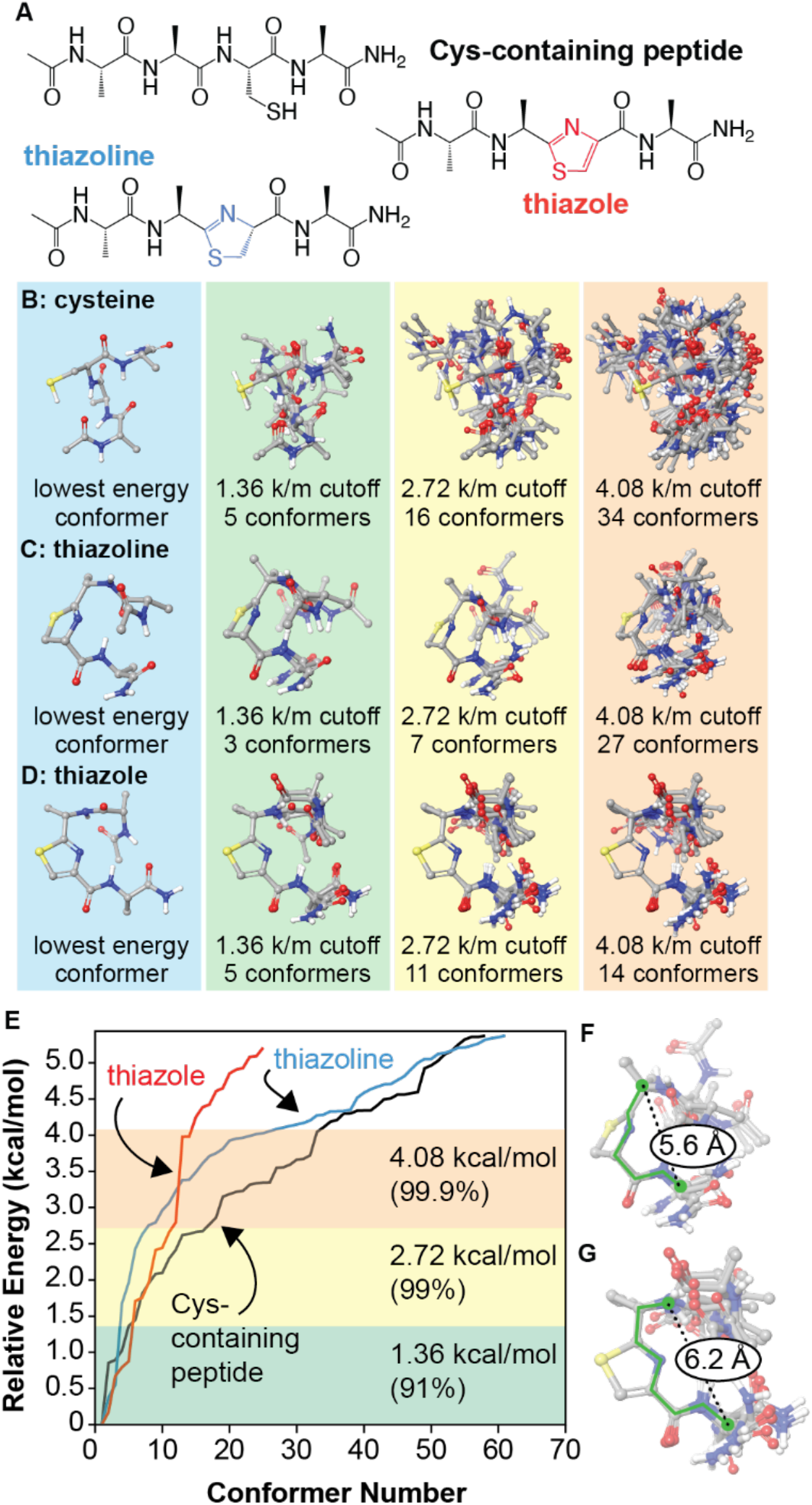
Conformational effects of thiazoline and thiazole formation.(A) The open chain and cyclized analogs Ac-AACA-NH_2_ were examined. Initial conformational searches were conducted using MacroModel (OPLS4 force field). All species within 4 kcal/mol of the global minimum were geometry optimized using DFT (B3LYP/6-31G**, SM8 solvent model) and re-ranked. (B-E) Lowest energy conformers are superimposed for different energy cutoff values. (F) A rigid 6-bond motif (green) describes all identified conformers within 2.72 kcal/mol of the global minimum for the thiazoline. (G) A similar 7-bond motif describes the thiazole conformers. k/m = kcal/mol. For images of all structures within 4.08 kcal/mol of each global minimum, see **Supplementary Materials**.

The results of the conformational analysis appear in **Figure 6B-E**, sorted by progressive energy cutoffs relative to each global minimum. The non-cyclized, cysteine-containing peptide exhibits the greatest flexibility, with 5 conformers being identified within 1.36 kcal/mol of the global minimum (91% of the population), and 16 within 2.72 kcal/mol (99%). Moreover, the identified conformers are largely non-superimposable, indicating that a high degree of conformational space is accessible within these energy ranges. In contrast, the thiazoline exhibits the most significant reduction in flexibility, with only 3 conformers identified at the 1.36 kcal/mol cutoff level and only 7 identified at a cut-off of 2.72 kcal/mol. Superposition of the thiazoline rings of these conformers reveals a rigid 6-bond motif that is preserved in all cases (**Figure 6F**). The thiazole analog exhibits similarly reduced flexibility, with 11 conformers being identified within 2.72 kcal/mol of the global minimum. In this case, a rigid 7-bond motif can be identified (**Figure 6G**). These evaluations provide the basis of models that could be used to predict the conformational effects of backbone cyclization on larger sequences and could be used to predict sequence locations in which cyclizations are more likely to be successful. In current experiments, we are combining molecular dynamics studies with experimental data to examine the longer-range effects that result from introducing thiazoline and thiazole groups in larger peptides and full-size proteins. Such information could be used to apply this chemistry more generally to improve the physio-chemical properties of therapeutic proteins.

### Conclusions

One can imagine two mutually synergistic strategies to introduce non-natural monomers into polypeptide and protein oligomers.^52^ One “bottom-up” approach relies on extant or engineered ribosomes to accept and process tRNAs carrying diverse non-canonical α-amino or non-α-amino acids.^53^ Hundreds of non-canonical α-amino acids (as well as α-hydroxy acids^54,55^) have been introduced into proteins in cells and animals using genetic code expansion,^56,57^ which usually relies on novel orthogonal aminoacyl tRNA synthetases to generate the requisite acylated tRNAs. Select non-canonical α-amino acids^58^ and one β-amino acid^30^ have also been incorporated into proteins *in vivo* using endogenous α-aminoacyl tRNA synthetases. Alternatively, many non-canonical α-amino acids, as well as certain non-α-amino acids, including β-amino acids^32,59^ and certain polyketide precursors,^33^ can be introduced into short peptides *in vitro* and on small scale using genetic code reprogramming, in which a stoichiometric RNA co-reagent (Flexizyme^60^) generates the requisite acylated tRNA.

The second “top-down” approach is reminiscent of late-stage functionalization reactions used to manipulate complex small molecule natural products^61,62^ and the natural biosynthetic strategy used to assemble ribosomally synthesized and post-translationally modified peptides (RiPPs).^13^ In this approach, enzymes, chemical reagents, or chemical catalysts are employed to post-translationally modify a peptide^12^ or protein^52^ to install a new or modified monomer. Examples of this approach include reactions of natural or non-canonical protein side chains or modification of the N- or C-terminus.^63–66^ The only backbone-focused non-enzymatic reaction of which we are aware is the *O*-mesitylenesulfonylhydroxylamine-promoted oxidative elimination of Cys residues to generate a dehydroalanine backbone^67^ that is subsequently modified. We note that the top-down and bottom-up strategies are complementary, and both have the potential to operate *in vivo* where very high protein titers are possible.^68^

Here we show that a constitutively active form of MicD and ArtGox, two enzymes used in the biosynthesis of cyanobactin natural products^69^ are sufficiently promiscuous to process substrates containing diverse backbone-modified monomers within substrate polypeptides, even at positions immediately preceding or following the site of cyclization/dehydrogenation. The backbone-modified monomers compatible with MicD-F and ArtGox include many accepted by extant ribosomes in small-scale *in vitro* reactions, including aramids and β^2^- and β^3^-amino acids. The products of these reactions are sequence-defined chemical polymers with multiple, diverse, non-α-amino acid monomers. We show further that cyclodehydration and dehydrogenation can install thiazoline or thiazole backbones within protein loops and linkers without disrupting the native tertiary fold. Calculations reported here reveal the extent to which these heterocycles restrict conformational space; they also eliminate a peptide bond–both features could improve the stability or add function to linker sequences now commonplace in emerging biotherapeutics. Moreover, as thiazoles and thiazoline heterocycles are replete in natural products,^5–7^ small molecule drugs,^8,9^ and peptide-mimetic therapeutics,^10^ their installation in protein-based biotherapeutics could improve or augment performance, activity, stability, and/or selectivity. More generally, this work represents a general strategy to expand the chemical diversity of the proteome without need for genetic manipulations.

## Acknowledgements

This work was supported by the NSF Center for Genetically Encoded Materials (C-GEM), CHE 2002182. We are grateful to Professor James Naismith (Oxford University) for graciously sharing plasmids and advice, and to Professor Susan Marqusee and for generous use of their CD instrument.

